# Sequential Amino Acid Mutagenesis-Driven De Novo Evolution of Adenine Deaminases Enables Efficient In Vivo Base Editing in Primate

**DOI:** 10.1101/2025.05.14.653640

**Authors:** Jiaoyang Liao, Hongling Zhang, Shuanghong Chen, Shenlin Hsiao, Chongping Lai, Hu Feng, Wendan Ren, Changrui Feng, Da Xie, Yanqiu Zheng, Wenhui Cai, Wenlong Wang, Yanhong Jiang, Dawei Wang, Erwei Zuo, Zijun Wang, Yuxuan Wu, Yuming Lu

## Abstract

Base editing allows for the precise modification of genetic information, providing new avenues for treating diseases^1^. The adenine base editor ABE8e is currently the most efficient and widely used tool for adenine base editing. ABE8e was developed through multiple rounds of directed evolution of Escherichia coli tRNA adenine deaminase, including phage-assisted continuous evolution (PACE)^2,3^. While PACE is highly effective, it is a complex system that poses challenges for implementation^4^. Despite its high efficiency, ABE8e is associated with limitations such as relatively higher bystander editing effects and elevated off-target activity^5^, which need to be addressed to further enhance its precision and safety. Here, we developed a novel method for the de novo discovery of evolved ABE components, particularly adenine deaminases. This process involves identifying candidate proteins through AI-based structural prediction and clustering, followed by the enhancement of deaminase editing activity through screening libraries created by sequential amino acid saturation mutagenesis. This evolutionary strategy simplifies the approach by employing saturation mutagenesis libraries tailored to specific segments, thereby enabling exploration of an expanded sequence space and increasing the likelihood of discovering adenine deaminases with superior capabilities. The newly developed hpABE5.20 here demonstrates a more refined editing window, reduced DNA off-target effects that are both sgRNA-dependent and -independent, and minimized RNA off-target activity, while maintaining robust editing efficiency relative to ABE8e. Furthermore, hpABE5.20 has been successfully applied for precise and effective therapeutic adenine base editing in cellular disease models, humanized mice, and non-human primates.

## Main

Theoretically, adenine deaminase can act on adenine (A) in DNA, converting it into inosine (I), which is subsequently interpreted as guanine (G) by the cell’s DNA replication machinery, thereby achieving an A-to-G base conversion^6^. High-precision genome targeting systems, such as CRISPR, can further direct adenine deamination to specific target sequences on the genome^7^. Researchers have combined these two approaches to achieve site-specific DNA base conversion^2^. The development of this system has introduced a novel technological advancement to the fields of genetic research and disease treatment. Specifically, adenine base editors (ABEs) were developed by fusing nCas9 with evolved TadA (eTadA), originally a tRNA adenine deaminase from *Escherichia coli*. This fusion enables efficient A•T to G•C conversions without significantly generating indels^2^.

The eighth-generation ABE, the state-of-art highly efficient ABE, was developed through multiple rounds of evolutionary optimization. Initially, the ABE prototype was refined through various molecular engineering approaches and bacterial screening-mediated directed protein evolution of tadA, leading to the development of ABE7.10^2^, an adenine base editor with enhanced editing efficiency. Building on this foundation, two research teams employed different directed protein evolution strategies to develop ABE8e^3^ and ABE8s^8^, which offer improved effectiveness and broader applicability. Among these, ABE8e was developed using phage-assisted continuous evolution. This method is highly automated but requires extensive expertise in microbial evolution and involves a complex system that is challenging to establish. In comparison, the bacterial screening-directed evolution used for ABE8s is relatively straightforward. However, the mutation depth of the customized random mutation library employed in this method is often constrained by the associated customization costs. Additionally, both strategies are limited in their ability to effectively simulate the co-evolutionary mutations involving multiple consecutive amino acids.

Here, to circumvent the shortcomings of current ABEs’ capabilities and commonly used evolution method, we employed a novel evolutionary strategy that involves using random primer amplification to construct a library of small-range continuous amino acid saturation mutations, followed by bacterial screening. The dominant variants identified through this screening are then integrated, either manually combination or as inputs for subsequent rounds of mutation screening. This strategy is akin to iterative saturation mutagenesis (ISM)^9,10^, but in this study, we broadened the scope and depth by employing a saturation mutation library covering up to four consecutive amino acids and conducting iterative saturation mutagenesis screening across the entire gene length. Our strategy emphasizes a thorough exploration of multiple consecutive amino acids, demonstrating ultra-high-throughput capabilities in protein sequence space exploration. This approach allows for the rapid screening of numerous variants, progressively evolving wild-type enzymes with minimal activity into highly active mutant deaminases, thereby facilitating the de novo development of base editing effectors.

In contrast to the development of ABE8e and ABE8s, which required multiple rounds of optimization—from the prototype to ABE7.10, and then from ABE7.10 to the eighth-generation ABE—we successfully developed an editor with efficiency comparable or even superior to ABE8e within a single study. More broadly, compared to the commonly used methods for constructing single-point or multi-point scattered mutation libraries^8,11-14^, our mutation strategy explored the sequence space of different amino acid combination mutation forms, achieved a higher-dimensional mutation space search, and resulted in a substantial enhancement of protein activity through highly successful directed evolution. As a result, our development strategy has led to the creation of several new ABEs, such as hpABE5.20, which exhibit editing efficiencies comparable to or even exceeding those of ABE8e, while also offering high precision. Notably, hpABE5.20 has demonstrated high-efficiency and high-precision therapeutic adenine base editing in disease cell models, human hematopoietic stem cells, and non-human primates. This work provides a promising approach that contribute to the advancement of base editing technologies and the development of more precise genetic therapies.

## Result

### Structural-clustering guided mining and de novo evolution of hpTadA

Following multiple rounds of directed protein evolution, the adenine base editor utilizing *E. coli* tRNA deaminase as the effector has demonstrated robust A-to-G base editing across various cell types^15-18^. Additionally, these ABEs have shown promising potential for both in vitro and in vivo gene editing applications^19-22^. However, the high off-target activity associated with its potent editing capabilities may present challenges for clinical use, necessitating further refinement of these base editors. Given these results, we sought to construct superior base editing tools with enhanced performances use deaminases newly discovered and evolved de novo.

There are four primary types of enzymes capable of deaminating adenine, represented by *E. coli* TadA^23^, human ADAT2^24^, human ADAR2^25^, and mouse ADA^26^. However, previous studies have shown that these natural adenine deaminases cannot accept DNA as a substrate and therefore cannot be directly used for adenine base editing (ABE)^2^. Given the successful development of a highly active adenine base editor from *E. coli* TadA, we used ecTadA as a starting point to identify new enzymes with potential functional capabilities based on structural similarity.

To this end, we initially compiled a set of 759 adenine deaminase candidate proteins (Table S1)., selected based on literature reports^27-29^, protein length, evolutionary relationships, and species representation. We then predicted their structures using AlphaFold2^30,31^ and clustered them using the TM-Cluster^32^ tool with a TM score threshold of 0.85 (Figure 1a,b). This process resulted in 22 clusters, ranging in size from 1 to 244 members (Figure S1). Notably, ecTadA is in cluster 1, and this cluster also includes 24 TadA proteins that exhibit high structural similarity to ecTadA (Figure S2) (Table S1). We synthesized the coding sequences of these proteins to replace TadA8e in ABE8e and assessed their deamination abilities in bacteria. Two of these proteins exhibited above-average editing activity (Figure 1c). We selected the protein with the highest natural activity(tadA005), TadA from *Hafnia paralvei*, for de novo evolution.

**Figure 1.**
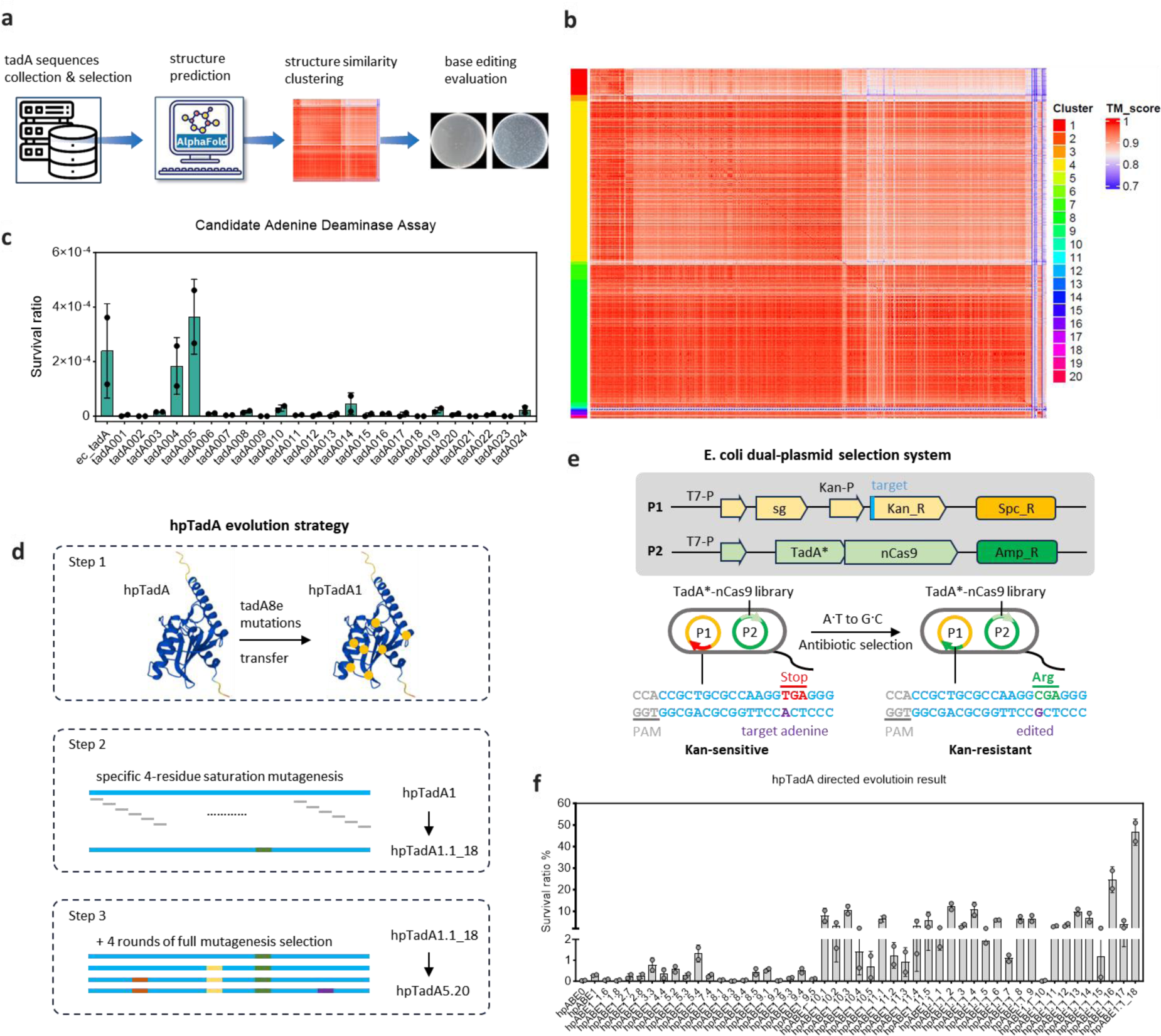
**a**, Schematic representation of TadA ortholog mining. Sequence collection and screening are followed by structure prediction, clustering, and deamination activity characterization. **b**, Heat map of structural similarity among 759 candidate TadA orthologs. Color blocks on the left indicate clusters identified using the TM-cluster method. **c**, Adenine deamination activities of 24 TadA orthologs in cluster 1, compared to Escherichia coli TadA. **d**, Evolutionary strategy of hpTadA: (1) Transfer TadA8e mutations to hpTadA; (2) Saturation mutagenesis of 11 specific 4-amino acid groups in the key region; (3) Four rounds of whole-gene mutagenesis screening. **e**, Schematic of the E. coli dual-plasmid screening system: One plasmid expresses sgRNA and contains the editing target at the 5’ end of the kanamycin resistance gene, creating a premature stop codon that inactivates the gene. The other plasmid expresses the adenine base editor. Effective A to G editing restores kanamycin resistance, enabling bacterial survival; otherwise, cells are eliminated. **f**, Directed evolution of hpTadA. Bacterial selection assays were used to evaluate the editing efficiency of ABE variants with different TadA mutations.

After selecting TadA005 (hereafter referred to as hpTadA), we first aligned its sequence with the highly efficient TadA8e variant from ABE8e. We then transplanted all 17 point mutations from TadA8e, relative to wild-type TadA (W21R, P46A, R49L, L82F, A104V, D106N, L107S, T109R, D117N, R120N, H121Y, S144C, F147Y, R150P, Q153V, K154F, and K155N), into hpTadA to create hpTadA1 (Figure 1d). In bacteria, the average editing efficiency of hpABE1, constructed using hpTadA1, reached 0.29% (Figure 1f), which is 7.95 times higher than the efficiency of hpABE0 (3.65 × 10⁻⁴), constructed with wild-type hpTadA. However, this also indicates that there remains significant room for further improvement.

To further enhance the deamination ability of hpTadA1, we designed a dual-plasmid bacterial screening system to screen variants generated through saturation mutagenesis of multiple consecutive amino acids (Figure 1e). In this system, one plasmid carries the elements for expressing the guide RNA, the spectinomycin resistance gene, and a kanamycin resistance gene expression cassette with a premature terminator as the editing target. The other plasmid harbors an ampicillin resistance gene and expresses base editors composed of high throughput-library of TadA variants. When a TadA variant is inactive or exhibits very low activity, the ABE it forms cannot efficiently deaminate the target adenine under the guidance of the guide RNA, leading to insufficient expression of the kanamycin resistance protein. Conversely, if the variant exhibits strong deamination activity, it will produce sufficient kanamycin resistance protein, allowing it to survive under the screening conditions and be selected (Figure 1e).

In protein engineering, different molecular evolution strategies enable the exploration of distinct protein sequence spaces, allowing for the development of adenine base editing tools that can surpass current ABEs in both efficiency and accuracy. Some protein functions may rely on the coordinated actions of multiple adjacent amino acids, where single mutations might not reveal these synergistic effects^9^. To address this, we employed a continuous amino acid saturation mutation strategy, diverging from the traditional single-point mutation approach, to conduct directed evolution across multiple consecutive amino acids. We identified 11 consecutive four-amino-acid (4aa) regions surrounding the 17 previously mentioned mutation sites and constructed 11 corresponding 4aa saturation mutation libraries (Figure1d). Through screening these libraries, we obtained several variants that exhibited notable improvements in editing efficiency during bacterial editing screening experiments. For instance, variants such as hpABE1_3.3 (average 0.77%), hpABE1_5.4 (average 1.33%), hpABE1_10.x (average 5.46%) from the 10th library, and hpABE1_11.x (average 4.31%) from the 11th library showed significant enhancements in editing performance (Figure 1f). Building on these results, we combined the mutations from hpABE1_3.3 and hpABE1_5.4 and used this combined variant as a template to construct libraries for further screening in region 10 and 11. Ultimately, this approach led to the development of the adenine base editor hpABE1_18, which achieved an average editing efficiency of 46.70% in bacteria (Figure 1f). This represents a 1280.4-fold increase compared to the 0.0365% average editing efficiency of the wild-type hpABE0.

### Enhanced A>G Editing with improved precision and reduced Off-Target Effects by Further **Evolved hpTadA**

Next, to evaluate the editing efficiency of hpABE1_18 in eukaryotic cells, we delivered its expression plasmid into HEK293T cells, targeting the PCSK9 site^19^ for installing therapeutic editing. We assessed its editing performance using NGS 48 hours post-transfection. At a lower transfection dose (100 ng/well in a 24-well plate) and without selection, the editing efficiency of hpABE1_18 in eukaryotic cells was found to be relatively low (4.46±0.14%) compared to ABE8e (32.84±4.01%) (Figure 2a). These initial results indicated that our regional mutation library approach is effective for generating improved mutants whereas much effort was needed to yield ABEs that convert A·T base pairs efficiently to G·C in human cells. Consequently, we proceeded with further bacterial screening and directed evolution of hpABE1_18. We implemented a continuous amino acid mutation strategy encompassing the entire gene segment and conducted an additional four rounds of mutation screening based on hpABE1_18. During these rounds, we gradually increased the screening intensity and varied the types of screening antibiotics (see Table S2 for screening conditions) to eliminate false positives that could arise from bacterial adaptive resistance to specific antibiotics. After each round of screening, all surviving variants were retained and pooled as mutation templates for the subsequent round, aiming to preserve as many different mutation branches as possible. Simultaneously, after each round, some variants were sampled to test their editing efficiency in HEK293T cells. Through these iterative rounds of evolution, the editing efficiency of ABEs constructed from these variants steadily improved, rising from an average of 13.90% in eukaryotic cells during the second round of evolution to an average of 28.41% by the fifth round (Figure 2a). In this final round of screening, several variants emerged with editing efficiencies comparable to or even surpassing those of ABE8e. We selected several mutants with high editing efficiency (hpABE5.13∼25) and tested their editing windows at two additional genomic sites containing multiple adenines within the editing window in HEK293T cells. It was found that while multiple variants maintained high editing efficiency, their editing windows were further narrowed (Figure 2b). Considering both efficiency and accuracy, we selected the best-performing variant, hpABE5.20, for subsequent research. To date, through a combination of structural clustering mining, bacterial screening, and directed evolution methods, we have developed an adenine base editor that demonstrates editing efficiency at the PCSK9 target site comparable to the most efficient and widely used adenine base editor, ABE8e, while offering greater precision (Figure 2b).

**Figure 2.**
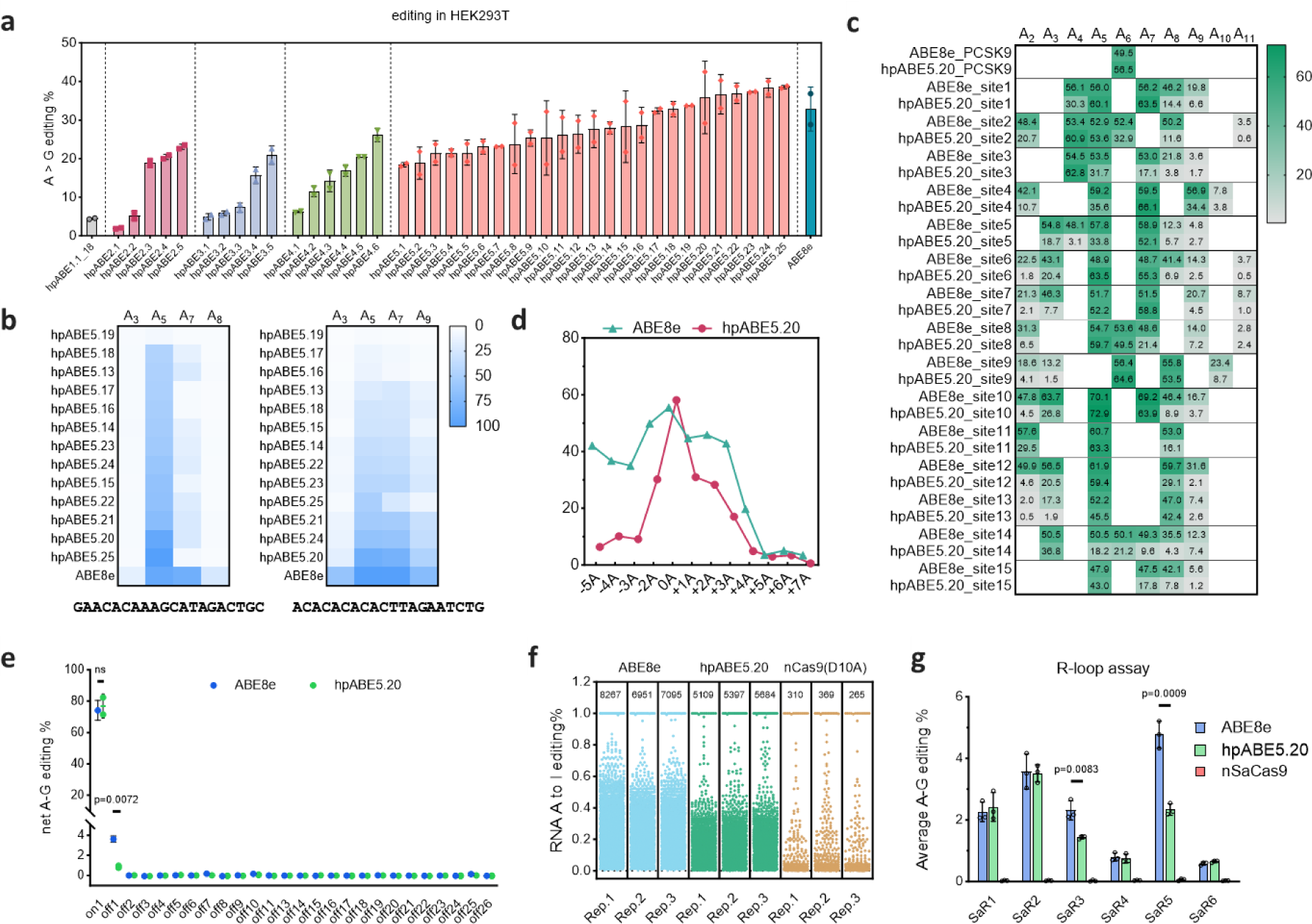
**a**, Editing efficiency of ABE composed of hpTadA variants transfected into hepG2 cells in the form of mRNA-LNP at PCSK9 site. Data are presented as mean±SD from two independent experiments. **b**, Editing precision of selected high-efficiency ABE variants at two additional sequences containing multiple adenine targets. **c**, Comparison of hpABE5.20 and ABE8e editing at 16 endogenous sites. Data are presented as the mean of three independent experiments. **d**, Comparison of editing precision between hpABE5.20 and ABE8e. Editing outcomes at 16 sites from panel **c** were analyzed, with 0A as the primary common substrate, and other positions defined relative to 0A. Data are presented as the mean of three independent experiments. **e**, Comparison of guide RNA-dependent DNA off-target editing between hpABE5.20 and ABE8e. Data are presented as mean±SD from three independent experiments. **f**, Comparison of RNA off-target editing between hpABE5.20 and ABE8e. Each point represents a detected RNA editing event. The total number of RNA A to I editing sites for each group is labeled at the top of the respective column. **g**, Comparison of guide RNA-independent DNA off-target editing at five sites between hpABE5.20 and ABE8e. Data are presented as mean±SD from three independent experiments.

For further characterization, we then conducted a comparison between hpABE5.20 and ABE8e across a broader range of endogenous loci. We assessed the editing efficiency and editing window of both editors at 16 endogenous sites^11^, including PCSK9 target site. Our results showed that at the vast majority of these sites (15 out of 16, with the exception of site 14), hpABE5.20 achieved editing efficiencies that were close to (>85%) or higher than those of ABE8e with substantially decreased bystander editing activity (Figure 2c). This demonstrates that hpABE5.20 provides superior editing accuracy than ABE8e at more sites. The positions of the primary editing target substrates at these 16 sites vary, and when we compared the editing windows based on the relative positions of the primary editing targets, it became clear that hpABE5.20 resulted in fewer bystander edits than ABE8e (Figure 2d). This finding confirms that hpABE5.20 not only matches or even surpasses ABE8e in efficiency but also offers enhanced editing precision.

Since off-target editing remains a major concern for therapeutic application, we then compared the off-target editing profiles of hpABE5.20 and ABE8e. To assess sgRNA-dependent DNA off-target editing, we first used Cas-OFFinder^33^ to predict 26 potential off-target sites harboring up to 3 bases mismatch with the human PCSK9-sg1 locus (Table S3). hpABE5.20 and ABE8e were subsequently transfected into primary human hepatocytes (PHH) using LNP-mRNA. Edited cells were collected, the editing sites and potential off-target sequences were amplified, and NGS was performed to quantify both on-target and off-target editing results^34^. The results indicated that the targeted editing efficiencies of ABE8e and hpABE5.20 were comparable, with ABE8e at 74.17 ± 4.53% and hpABE5.20 at 76.89 ± 5.46%. Both editors exhibited detectable off-target editing at the off1 site, but hpABE5.20 showed significantly lower off-target activity at this site (0.87 ± 0.10%) compared to ABE8e (3.62 ± 0.21%), representing approximately one-fourth of the off-target activity observed with ABE8e (Figure 2e). In addition to DNA off-target effects, adenine base editors can cause A-to-I RNA editing within cells^35^. To investigate this, we transfected hpABE5.20 and ABE8e into 293T cells using PEI, delivering the editors in plasmid form. Cells were collected 48 hours post-transfection, and transfection-positive cells were enriched via flow cytometry. RNA was extracted, and transcriptome sequencing was performed to identify A-to-I editing events across the transcriptome. Compared to the control group (nCas9), both hpABE5.20 and ABE8e induced A-to-I RNA editing. However, the number of RNA off-target editing events caused by hpABE5.20 was significantly lower than that caused by ABE8e, with a reduction from an average of 7,437 to 5,396 off-target events (p=0.0104) (Figure 2f). The fused TadA in ABEs may also deaminate DNA in the unwinding region of the cell in an sgRNA-independent manner^3^, potentially leading to sgRNA-independent DNA off-target effects. To detect this form of off-target activity for hpABE5.20 and ABE8e, we utilized nSaCas9 to create R-loops at six specific sites within the cell and then assessed the editing effects of hpABE5.20 and ABE8e at these sites. After transfection and screening, the genomes of the treated cells were extracted and the sequence encompassing tested sites were amplified for NGS to detect the frequency of off-target editing. We found that four of the sites showed similar off-target editing frequencies between hpABE5.20 and ABE8e. However, at two sites, the off-target editing caused by hpABE5.20 was significantly lower (p < 0.01) compared to ABE8e (Figure 2g). These findings collectively suggest that hpABE5.20 not only matches ABE8e in on-target editing efficiency but also offers improved specificity by reducing off-target effects in both DNA and RNA, making it a potentially safer option for clinical applications.

### Comprehensive Unbiased Target Analysis of hpABE5.20 Using Paired sgRNA-Target Libraries

We next sought to comprehensively and impartially analyze the editing performance of hpABE5.20 and ABE8e. We designed an sgRNA-target library containing 20,480 pairs (Figure 3a). These library fragments were first synthesized and then cloned into vectors to create lentiviral libraries, which were used to infect 293T cells to establish stably transfected target cell lines. Subsequently, we transfected these cells with ABE plasmids, and high-throughput sequencing was then performed to quantify the editing outcomes (Figure 3a). Intriguingly, we analyzed the editing efficiencies across all possible single-nucleotide flanking base compositions and found that hpABE5.20 exhibited higher average editing efficiencies in the contexts of CAT, CAC, TAC, and TAT (Figure 3b). Similarly, when examining all possible two-base compositions, we observed that hpABE5.20 preferentially edited the adenine in the CA and TA motifs over the AA and GA motifs, and the average editing efficiency for AC and AT was also higher than for AA and AG (Figure 3b). Direct analysis of sequence preferences at the editing site revealed that, compared to ABE8e, which slightly favors YA sequences (consistent with previous research findings^36^), hpABE5.20 shows a slight preference for YAY sequences (Figure 3c). This preference may be related to the fact that the target substrate used during the evolutionary process was CAC (Figure 1e). These results implied that by adjusting certain sequence condition harboring targeting adenines use in the screening system, out evolution strategy might yield evolved deaminase preferentially for specific motif sequence and thus more applicable for therapeutic edits under certain sequence context. We then compared the editing window of hpABE5.20 and ABE8e across all target sites. Results from two independent experiments indicated that, compared to ABE8e, whose primary (>80%) editing window was concentrated at the A4-A7 positions, the main editing window of hpABE5.20 was more focused on the A6 and A7 positions (Figure 3d). This suggests that hpABE5.20 not only maintains high editing efficiency but also offers enhanced precision by narrowing the editing window.

**Figure 3.**
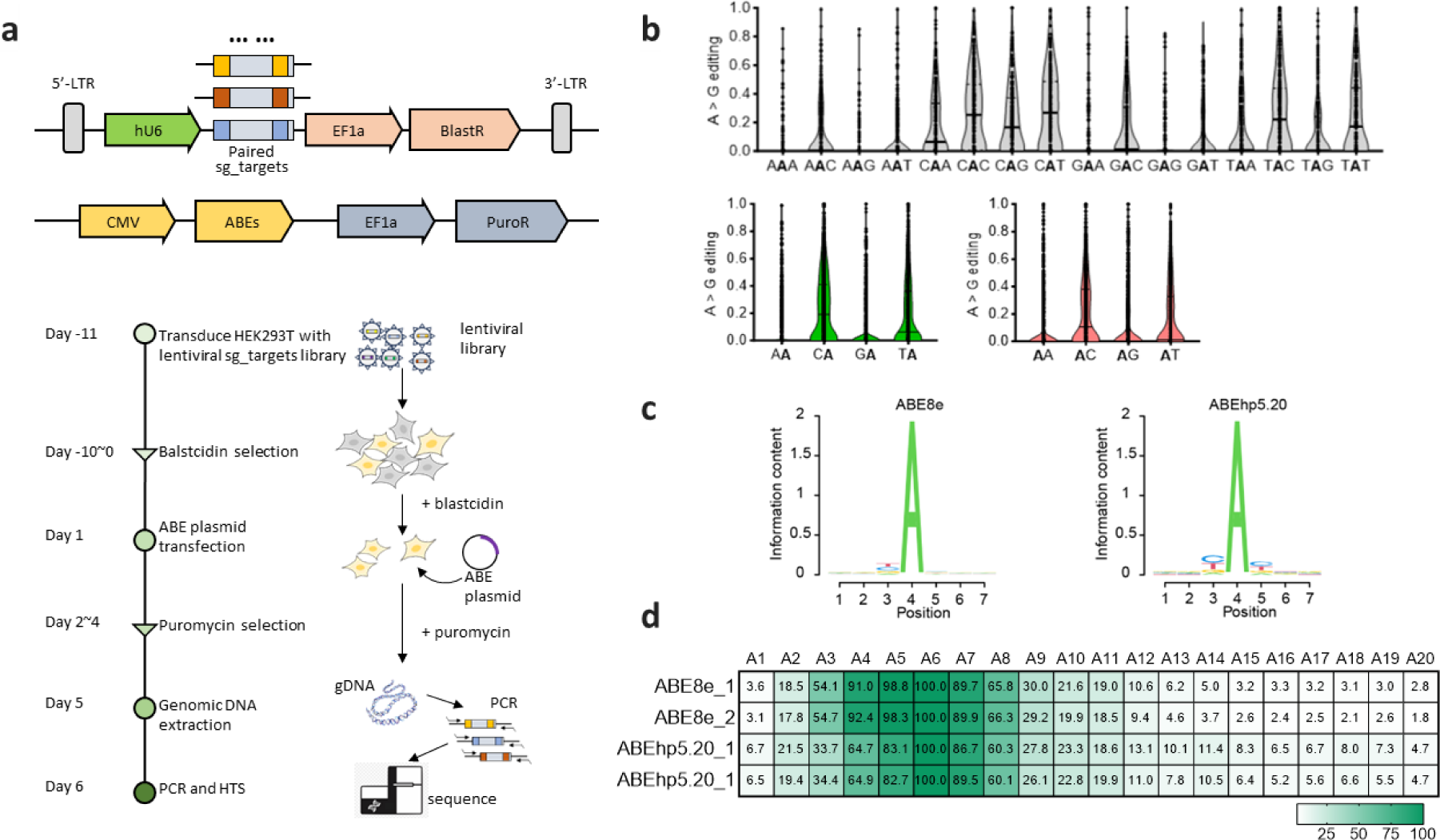
**a**, Schematic of the experimental workflow for characterizing ABE editing performance using a library. Paired sg_targets are introduced into cells via lentiviral transduction to create stable cell lines. ABE expression plasmids are then transfected into these cells using PEI for base editing. Successfully transfected cells are selected with antibiotics, followed by collection, genomic DNA extraction, target sequence amplification, and quantification of editing results via high-throughput sequencing. **b**, Analysis of target adenine editing across different contexts. Bolded adenines indicate the target sites. Editing efficiencies of all target sites are visualized using violin plots. The top row lists all possible three-base compositions, while the bottom row shows two-base compositions. Results are from two independent experiments with similar outcomes. **c**, Weblogo representations of the sequence preferences for the substrate contexts of ABE8e and hpABE5.20. The experiments were conducted twice with consistent outcomes. **d**, Comparative analysis of the relative A to G editing efficiency at various positions for ABE8e and hpABE5.20. Results are presented from two independent experimental replicates.

### hpABE5.20 supports robust editing in various of disease-relevant context with particular success in the in vivo base editing of non-human primates

To explore the therapeutic potential of hpABE5.20 for treating genetic diseases, we selected the PAH gene mutation as a model and assessed its ability to correct two pathogenic missense mutations: PAH_c.721C>T (p.Arg241Cys) and PAH_c.1243G>A (p.Asp415Asn) for which single-nucleotide editing with improved precision is critical^37^. There are multiple potential adenines editing sites at positions 4-13 within the spacers targeting these missense mutations. The ABE expression plasmid was transfected into two stable cell model lines harboring the aforementioned pathogenic mutations. At the PAH_c.721C>T site, ABE8e not only corrected the pathogenic mutation with an average efficiency of 50.03% but also exhibited high editing activity at positions 7 and 8 (average A7 at 45.22% and A8 at 27.27%). In contrast, hpABE5.20 demonstrated high-efficiency editing primarily at the pathogenic site, with an average of 44.63% targeted editing, while undesired bystander editing at A7 and A8 was significantly lower (11.11% and 4.24%, respectively) (Figure 4a). The average editing specificity at this site with hpABE5.20 was 2.59 times greater than that of ABE8e (Figure 4c). Similarly, at the PAH_c.1243G>A site, hpABE5.20 also exhibited higher editing precision compared to ABE8e, with a specificity that was 1.84 times greater (Figure 4b,c). These results suggest that while hpABE5.20 is as efficient as ABE8e in correcting disease-causing mutations, and it offers significantly greater precision, reducing unwanted bystander editing in cellular models. This increased specificity makes hpABE5.20 a promising candidate for therapeutic applications in genetic disease treatment.

**Figure 4.**
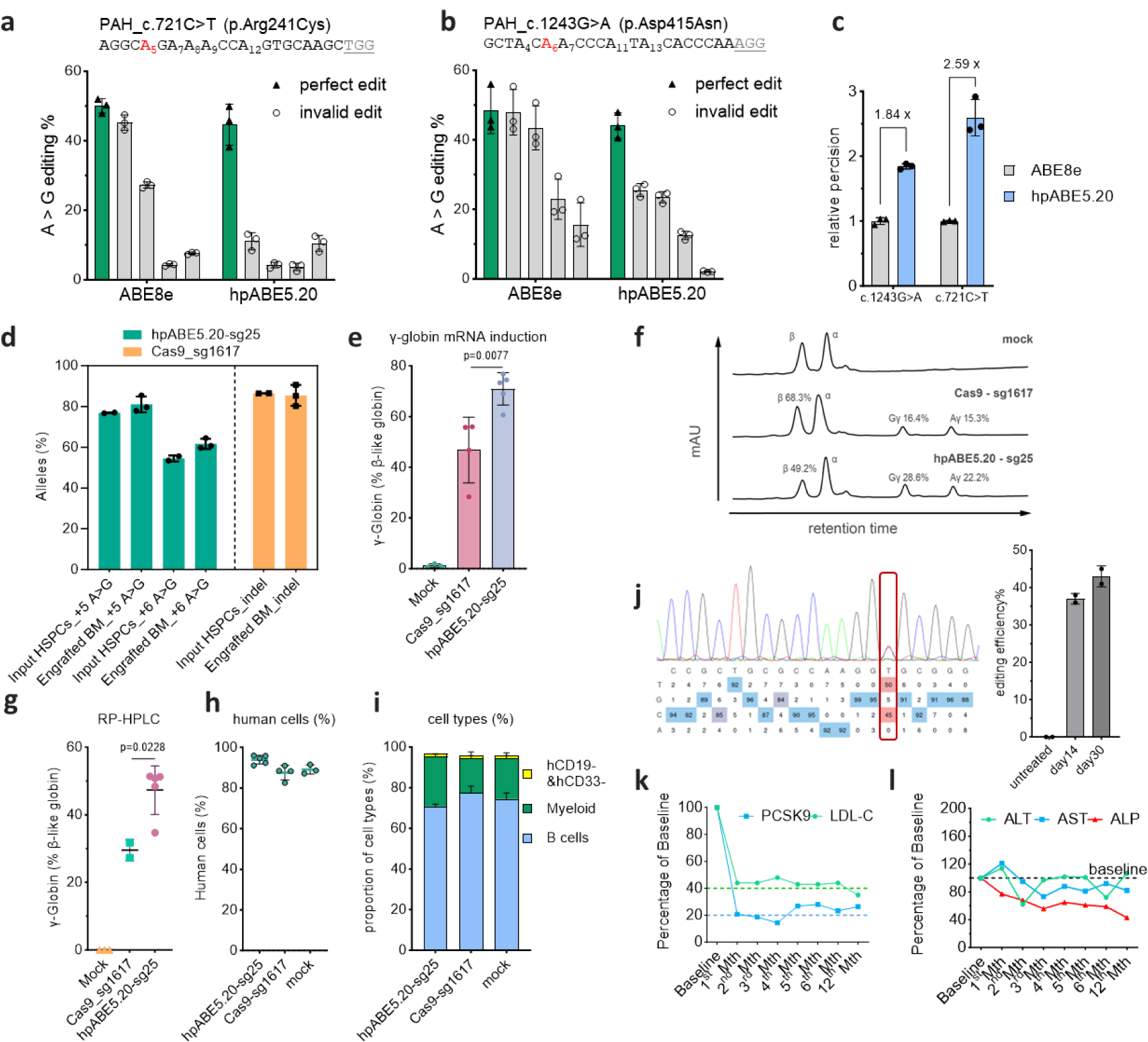
**a**, Comparative editing outcomes of ABE8e and hpABE5.20 on the PAH_c.721C>T mutation in a cellular model. Results are depicted as the mean±SD from three independent experiments. **b**, Comparative editing outcomes of ABE8e and hpABE5.20 on the PAH_c.1243G>A mutation in a cellular model. Results are depicted as the mean±SD from three independent experiments. **c**, Quantitative comparison of the relative precision of ABE8e and hpABE5.20 in repairing the cellular models. Data are presented as the mean±SD from three independent experiments. **d**, Base editing in unfractionated BM after 16 weeks as compared to input HSPCs. Data are plotted as mean ± sd. *n* =2 for input HSPCs, *n* = 5 primary recipients for each group of edited HSPCs. **e**, γ-globin induction analyzed by RT-qPCR normalized by β-like globin, measured from BM chimerism 16 weeks after base edited HSPC infusion. Data are plotted as mean ± sd. *n* = 5 replicates from individual recipient mice. **f**, γ-globin induction analyzed by RP-HPLC normalized by β-like globin, measured from BM chimerism 16 weeks after base edited HSPC infusion. Data are plotted as mean ± sd. *n* = 5 replicates from individual recipient mice. **g**, representative RP-HPLC results from **f**. **h**, Comparing human CD235a+ cells of mock with all edited groups. Data are plotted as mean ± sd. *n* = 5 mice from mock and all edited groups. **i**, Comparing human chimerism of mock with all edited groups. Data are plotted as mean ± sd. *n* = 5 mice from mock and all edited groups. **j**, Percentage of engrafted human B cells, myeloid cells and CD19^−^CD33^−^ cells 16 weeks after transplantation. Data were presented as mean ± sd, *n* = 5 mice. **k**, Temporal dynamics of PCSK9 expression and LDL-C levels in primates following in vivo base editing with hpABE5.20 & PCSK9-sg1 via LNP-mRNA delivery. **l**, Temporal dynamics of ALT, AST, ALP levels in primates following in vivo base editing with hpABE5.20 & PCSK9-sg1 via LNP-mRNA delivery.

Previously we and others have shown that Cas-mediated gene editing in BCL11A or HBG of hematopoietic stem cells (HSCs) to produce durable fetal hemoglobin is a practicable therapeutic strategy to ameliorate the server beta-globin disorders sickle cell disease and transfusion-dependent beta-thalassemia^38-40^. In contrast to Cas9, base editing is a genome editing method that directly generates precise point mutations in genomic DNA without causing extensive DNA double-strand breaks (DSBs) which might result in undesired indels, translocations and rearrangements. Next, we examined the editing efficiency of hpABE5.20 in human regenerative HSCs. Mobilized CD34+ HSPCs from a healthy donor were treated with various gene editing RNP complexes, including hpABE5.20-sg25 (AGGCAAGGCTGGCCAACCCA<u>TGG</u>, targeting HBG) and Cas9-sg1617 (CTAACAGTTGCTTTTATCAC<u>AGG</u>, targeting BCL11A). The sg25 target is an adenine base editing site identified in an unpublished study, which has been shown to efficiently induce gamma-globin expression. The Cas9-sg1617 control utilizes the same editing strategy as we previously tested^38,41^ and the currently commercialized Casgevy^39^. Following treatment, the edited and untreated cells were injected into NCG-X (NOD-Prkdc^em26Cd52^ Il2rg^em26Cd22^ Kit^em1^Cin(V831M)/Gpt) immunodeficient mice^15^ via the retinal venous plexus injection method. After transplantation, we assessed the editing efficiency and reconstitution of the human hematopoietic system in bone marrow (BM) samples isolated post-transplantation. The overall editing frequency showed similar outcomes in both the input and transplanted human cells (Figure 4d). RT-qPCR analysis of unfractionated BM cells revealed that gamma-globin mRNA levels were significantly increased in the edited groups compared to unedited control samples, accounting for 1.18% of total beta-like globin transcripts in the unedited control, 46.73% with Cas9-sg1617, and 70.88% with hpABE5.20-sg25 (Figure 4e). Furthermore, RP-HPLC analysis (Figures 4f, g) demonstrated that in erythroid cells (CD235a+ cells) differentiated in vivo following the editing and transplantation of HSPCs, the hpABE5.20-sg25 editing strategy induced significantly higher levels of γ-globin compared to the Cas9-sg1617 strategy, with an average of 47.32% versus 29.52% of total β-like globin (p=0.0228).

Flow cytometry analysis using anti-human and anti-mouse CD45 antibodies showed that human cells constituted over 85% of the BM in all mice, with an average of 89.09% in the control group, 93.98% in the hpABE5.20-sg25 group, and 87.56% in the Cas9-sg1617 group (Figure 4h). Additionally, flow cytometry quantified the relative abundance of human myeloid (hCD33+), B lymphoid (hCD19+), and hCD19−/hCD33− cells, revealing that the proportions of each lineage were approximately equal in mice receiving either unedited or edited cells (Figure 4i). This indicates that hpABE5.20-sg25-treated HSPCs maintained their capacity for engraftment and multilineage differentiation. Collectively, these findings demonstrate that base editing with hpABE5.20-sg25 does not compromise the engraftment or pluripotency of transplanted HSCs, while more efficiently inducing gamma-globin expression compared to currently commercialized editing protocols. This positions hpABE5.20-sg25 as a potentially superior tool for therapeutic gene editing in regenerative medicine.

In preparation for the clinical development of hpABE5.20, we used the PCSK9 target (CCCGCACCTTGGCGCAGCGG) as a model to evaluate the potential of hpABE5.20 for in vivo base editing in non-human primates. Targeted base editing of the PCSK9 gene can inhibit its expression, reduce LDL-C (Low-Density Lipoprotein Cholesterol) levels, and potentially provide a treatment for familial hypercholesterolemia^19^. To facilitate this study, we developed a novel, patented LNP-mRNA delivery system (Patent ID: CN 114989182 B) specifically designed for base editing studies in animals. We administered a single intravenous dose of 2 mg/kg to the experimental monkeys. The results demonstrated that hpABE5.20 achieved efficient base editing in vivo. At 14-and 30-days post-administration, the average editing efficiencies in right liver lobe biopsy tissues were 37% and 43%, respectively (Figure 4j). Prior to administration, the PCSK9 protein level was 47.8 ng/ml. Within the first month post-administration, the protein level dropped to 9.9 ng/ml, representing a 79.3% decrease. Notably, 15 months after administration, the PCSK9 protein level remained stable at around 12.3 ng/ml, a level sufficient to effectively treat hypercholesterolemia. Additionally, direct testing of LDL-C content in the blood showed a significant reduction within one month after administration, with levels decreasing from 1.325 mmol/L to 0.59 mmol/L, and remaining stable for 15 months. After 15 months, LDL-C levels further decreased to 0.5 mmol/L, indicating a sustained and effective therapeutic outcome. Throughout the study, we also monitored the levels of ALT (Alanine Aminotransferase), AST (Aspartate Aminotransferase), and ALP (Alkaline Phosphatase) in the blood of the experimental animals at various time points before and after administration. The results showed that these indicators remained stable or slightly decreased, suggesting that our in vivo base editing system has a favorable safety profile. These findings demonstrate that hpABE5.20, delivered via our patented LNP-mRNA system, not only achieves efficient and sustained base editing in non-human primates but also provides a significant and lasting therapeutic effect with a favorable safety profile, supporting its potential for clinical development.

## Discussion

In this study, we employed a structural similarity clustering method to discover the naturally active hpTadA and utilized directed evolution, with a focus on saturation mutation library screening of several consecutive amino acids, to develop the initially low-activity hpABE0. This variant was subsequently engineered into a new adenine base editor, hpABE5.20, which surpasses ABE8e in both editing efficiency and accuracy. Compared to ABE8e, hpABE5.20 exhibits lower sgRNA-dependent and -independent DNA off-target activity, as well as reduced RNA off-target editing. Furthermore, hpABE5.20 demonstrates efficient and precise A>G therapeutic editing capabilities in cell models, humanized mice, and non-human primates, underscoring its strong potential for therapeutic applications.

The development of hpABE5.20 has significantly enriched the base editing toolbox. With rational design or additional rounds of optimization and modification, its accuracy could be further enhanced, such as by narrowing the editing window and further reducing off-target effects. Moreover, by adjusting the editing target and selecting criteria, the evolution system we installed might yield ABE variants preferentially editing adenine under certain sequence conditions and thus more applicable for specific therapeutic scene.Additionally, it is feasible to develop other types of base editing systems by altering the base type of the primary editing product or substrate.

Structural similarity may better correlate with the functional similarity of proteins than sequence similarity. Recently, several paradigm-shifting studies in gene editing have focused on developing new editing tools based on structural similarities^32,42,43^. In one of our previous studies, we found that the TM cluster was exceptionally effective in situations where the average structural similarity between proteins was high^32^. Similarly, in this study, the TM cluster effectively facilitated the discovery of naturally active adenine deaminases, demonstrating its potential applicability in the structural mining of other functional proteins.

We employed multiple rounds of sequential amino acid saturation mutagenesis to explore a broader range of protein sequence space, thereby expanding diversity and increasing the likelihood of obtaining advantageous variants. Our research suggests that high-depth exploration of protein sequence space is a viable approach to significantly enhancing the activity of adenine deaminase on DNA, akin to “fine-tuning” specific regions of the protein. However, the general applicability of this method for engineering other enzymes still requires further research.

Additionally, our results demonstrated that delivering hpABE5.20-sg25 editing components to HSCs from healthy donors in the form of RNPs, followed by transplantation into immunodeficient mice, effectively induced gamma-globin expression. This supports the potential of hpABE5.20-based base editing as a new treatment option for β-hemoglobinopathies. Furthermore, hpABE5.20-sgPCSK9 delivered as LNP-mRNA successfully achieved safe and effective in vivo base editing in non-human primates. These findings highlight the significant application potential of hpABE5.20 developed in this study.

Overrall, our work established a rational evolution strategy providing a pragdigim for protein evolution and yeilded ABE variants with roubust editing efficiency, greatly expanding the toolchest for theraputic genome editing.

## Methods

### Adenine Deaminase Discovery and Phylogenetic Tree Construction

To discover new adenine deaminases, we downloaded microbial metagenomic data from the NCBI Sequence Read Archive (SRA) as our search database (https://www.ncbi.nlm.nih.gov/sra). We utilized HMMER3 to build a hidden Markov model (HMM profile) of TadA. Using hmmsearch with default parameters, we queried the database. After filtering for completeness and sequence length, we identified a total of 1028 TadA sequences. These were further processed for redundancy reduction using cd-hit. Subsequently, we employed iqtree with default parameters to analyze phylogenetic relationships relative to the reference TadA, selecting 20 sequences for further investigation and base editor construction to assess their adenine base editing activity. Ultimately, this process led to the identification of 9 new deaminases demonstrating adenine base editing capability.

### Plasmid Construction

The plasmid for expressing adenine base editors in bacteria is based on the J23119 vector (gift from Professor Yumin Lu), with CMV promoter-driven nCas9 (D10A) fused to various excavated or evolved deaminases and their variants. These deaminases were constructed into the vector by PCR (KOD-plus-neo, or Takara PrimerStar) followed by Gibbon assembly. PCR products were recovered using an agarose gel DNA recovery kit (Tiangen, DP209) and assembled using Gibbon recombinase (Vazyme ClonExpress II One Step Cloning Kit, C112). Other fragment elements that need to be assembled were also constructed into the target vector by Gibbon assembly, unless otherwise specified. The editing target was introduced into the ccdb toxic gene on the pBAD vector (gift from Professor Yumin Lu) by mutant primer PCR followed by Gibbon assembly. The adenine base editor plasmid for eukaryotic cell experiments was also constructed by PCR and Gibbon assembly based on ABE8e (Addgene, #138489). The plasmid vector expressing sgRNA (Addgene, #52963) was digested with Esp3I (ThermoFisher, FD0454) and recovered with a recovery kit (TIANGEN General DNA Purification Kit, DP214). Oligo forward and reverse primers were heated at 95°C in a PCR instrument for 2 minutes and then gradually cooled to anneal to form double-stranded fragments. The digested plasmid backbone and annealed fragments were reacted with DNA ligase (New England BioLabs, M0202) at room temperature for 30 minutes. The cloned plasmid was transformed into chemically competent cells DH5α (TransGen, CD201) and selected with the corresponding antibiotics. The plasmid for transfection was obtained using an endotoxin-free plasmid extraction kit (TIANGEN, DP117 or DP118).

### Directed evolution and efficiency evaluation of TadA005

We used a bacterial selection system to perform directed evolution to improve the editing activity of TadA005. The system consists of two plasmids, one of which is a bacterial expression vector (Amp_R) containing an adenine base editor fused to a TadA005 variant with nCas9(D10A). The vector also contains an expression frame for a guide RNA that can target the complementary sequence of the protospacer adjacent motif (PAM) in the ccdB toxin gene. The other plasmid (Cam_R/Kan_R) contains the ccdB gene regulated by the arabinose promoter pBAD. The editing target is located in the ccdB toxic gene on the pBAD vector. The vector will express the toxic gene to kill cells in the medium supplemented with L-arabinose, while the successfully edited cells will terminate the expression of the ccdB toxic gene prematurely and survive to achieve positive selection.

To generate a mutant library, mutations were introduced at specific positions of TadA005 using degenerate base primer PCR. Specifically, the single-side primer covers the amino acid position to be mutated and its flanking sequence position, and replaces the amino acid position to be mutated with multiple Ns. For example, if the library is 6 amino acid mutations, 18N is introduced. BsaI restriction sites should be introduced at both ends of the primer so that after completing PCR, they can be assembled using the Golden Gate assembled kit. The deaminase has a total length of 169 amino acids. Generally, a library is constructed for every 6 amino acids, and 28 libraries are constructed per round of evolution. The plasmid library of each round is electrotransformed into the Top10 electrotransformation competent medium that also contains the target plasmid, grown and screened on the corresponding resistance plate, and the grown clones are collected as the template plasmid for the next round of library construction, and 28 libraries are constructed again for screening. In this study, 5 rounds of selection were performed. In order to reduce the impact of acquired antibiotic adaptation from non-mutant library sources that may be produced by bacteria on the screening, and to gradually increase the screening intensity to obtain more active deaminase variants, we used different types and concentrations of antibiotics in 5 rounds of selection. Specifically, chloramphenicol (Cam) was used in the first and second rounds at concentrations of 4, 8, 16 µg/mL and 16, 32, 64 µg/mL, respectively. Kanamycin (Kan) was used in the third to fifth rounds at concentrations of 8, 16, 32 µg/mL, 32, 64, 128 µg/mL, and 128, 256, 384 µg/mL, respectively.

To evaluate the editing efficiency in bacteria, the electroporated cells were recovered in 1 mL of SOB medium at 37°C for 60 minutes. Subsequently, 5 µL of the bacterial culture was inoculated onto a control plate containing only Cam/Kan. The remaining culture was concentrated and inoculated onto another plate containing arabinose along with Cam/Kan. The editing efficiency was determined by calculating the percentage of CFUs in the induction plate relative to the control plate, based on equal volumes of bacterial suspension.

### Plasmid Transfection and Editing Efficiency Assessment

HEK293T cells were cultured in Dulbecco’s Modified Eagle Medium (DMEM) (Gibco, C11995500BT) supplemented with 10% fetal bovine serum (FBS) (Gibco, 10099141) and 1% penicillin/streptomycin (BasalMedia, S110JV) at 37°C in a humidified atmosphere with 5% CO_2_. HEK293T cells (50,000-100,000) were seeded in 48-well plates (Corning) approximately 24 hours prior to transfection to achieve 60-70% confluency. For transfection, 250 ng of base editor plasmid and 250 ng of sgRNA expression plasmid were co-transfected into the cells using PEI (Polysciences) following the manufacturer’s instructions. After 24 hours, selection was initiated using medium containing 2 µg/mL puromycin for 48 hours. Cells were harvested 96 hours post-transfection, and genomic DNA was extracted using the TIANGEN Genomic DNA Extraction Kit (DP304) or cell lysates were prepared using the Direct PCR Kit (Selleckchem, B40013). The target genomic region was PCR amplified using KOD-Plus-Neo (TOYOBO, KOD-401), and the amplicons were sequenced with 2x150 paired-end reads on the MiSeq Sequencing System (Illumina). Editing efficiency was analyzed using the default parameters of CRISPResso2 software.

### Construction of PAH Cell Model

First, a DNA fragment containing two ESP3I restriction sites in opposite orientations was inserted into the lentiviral vector lenti-blast using the previously described PCR-Gibson assembly method. Five pairs of oligos containing PAH disease point mutations (spacer + PAM 23nt, with 6nt upstream and downstream) were synthesized and annealed, then inserted into the lenti-blast vector through restriction enzyme ligation. This transfer plasmid was co-transfected into 293T cells along with the packaging plasmid psPAX2 and the envelope plasmid pMD2.G to produce lentiviral particles. The viral particles were collected by centrifugation and concentration, and their titer was determined before infecting 293T cells. Following infection, cells underwent selection with blasticidin at a concentration of 10 µg/mL for 10 days to establish stable cell lines.

### Library Analysis of Editing Characteristics of Adenine Base Editors

A plasmid library containing spacers and their corresponding target sequences was constructed. Each spacer was 20 nucleotides (nt) in length, with the fourth, fifth, sixth, seventh, and eighth positions fixed as adenine (A) and exhaustive variation in the surrounding ±3 nt positions, generating 4096 unique sequences per position, totaling 20480 spacers. For each spacer, the target sequence comprised a random 6 nt segment from the human genome, followed by the spacer sequence, a 3’ NGG protospacer adjacent motif (PAM), and another random 6 nt segment from the human genome. The library design included 20 bp overlapping sequences flanking both ends for incorporation into the lentiGuide-Puro vector (Plasmid #52963) driven by a human U6 promoter. This library was synthesized by Agilent (Agilent SurePrint Oligonucleotide Library Synthesis platform, G7238A), and assembled into the vector using Gibson Assembly.

The resulting plasmid library was packaged into lentiviral particles by transfecting 3 × 10^6 HEK293T cells seeded in 10 cm dishes with a total of 20 µg plasmid DNA using 80 µl polyethylenimine (PEI) (Polysciences, 23966). This included 10 µg plasmid expressing the sgRNA library, 6.6 µg lentiviral packaging plasmid, and 3.4 µg envelope plasmid. Medium was replaced 6 hours post-transfection, and viral particles were collected and concentrated 48 hours post-transfection. Viral titers were determined by serial dilution combined with antibiotic selection.

For lentiviral transduction, HEK293T cells were infected at a multiplicity of infection (MOI) of 0.3, with medium change 6 hours post-infection. To generate stable cell lines expressing the 20480 sgRNA-target library, cells were selected with 2 µg ml^-1 puromycin (Beyotime, ST551) for 7 days.

For evaluating base editor performance, 6 × 10^6 HEK293T cells stably expressing the library were seeded in 10 cm dishes and transfected with 20 µg plasmid encoding the base editor using 80 µl PEI at 70% confluency. Genomic DNA (gDNA) was extracted 72 hours post-transfection (Tiangen, DP304-03), and target regions were amplified for deep sequencing. The results of contextual preference motif analysis of adenine editing were produced using weblogo (https://github.com/WebLogo/weblogo).

### Detection of RNA Editing

To evaluate guide RNA-independent off-target RNA editing by the base editor, HEK293T cells were co-transfected with plasmids encoding both the base editor, EGFP and the sgRNA. Seventy-two hours post-transfection, the top 15% EGFP-positive cells were sorted by flow cytometry. The sorted cells were centrifuged, resuspended in 1 ml Trizol (Ambion, 15596018), and then shipped on dry ice to Personalbio (Shanghai) for mRNA extraction, library construction, and sequencing analysis. Paired-end 2×150-bp sequencing was performed on the Illumina platform, generating approximately 50 million reads per sample. Raw sequencing data were processed to remove adapters and low-quality reads. Specifically, 3’ end adapter sequences were trimmed using Fastp(https://github.com/OpenGene/fastp), and reads with an average quality score below Q20 were discarded. Filtered reads were aligned to the reference genome (Ensembl GRCh38) using HISAT2 (https://daehwankimlab.github.io/hisat2/). SNPs were called using Varscan (https://github.com/dkoboldt/varscan) with the following criteria: base quality score >20, read depth >8 at the SNP site, ≥2 reads supporting the variant allele, and SNP p-value <0.01. Variant loci of ABE8e, ABE8.8 or yolBE3.1 overexpression were filtered to exclude sites without high-confidence reference genotype calls of the control experiment, to ensure that the remaining variant sites are caused by the specific editing of the base editor. These sites were required to have at least 99% of reads containing the reference allele in control samples. RNA edits were further filtered to include only sites with >20 reads and at least one read containing the variant allele. A-to-I edits on the positive strand and T-to-C edits on the negative strand were both considered as A-to-I base edits.

### sgRNA-dependent DNA Off-Target Analysis

Potential off-target sites for PCSK9 sgRNA were predicted using Cas-OFFinder software(http://www.rgenome.net/cas-offinder/), with a maximum of 3 mismatches, 1 DNA bulge, and 1 RNA bulge. This analysis identified 54 potential off-target sites, which were reduced to 27 unique sites after removing duplicates. In vitro transcribed mRNA and chemically synthesized sgRNA were electroporated into HEK293T cells using the Lonza SF Cell Line 4D Nucleofector (V4XC-2032). Genomic DNA was extracted 96 hours post-electroporation using the Blood/Cultured Cell/Tissue Genomic DNA Extraction Kit (Tiagen, DP304). Primers for amplifying the potential off-target sites were designed using Primer3(https://primer3.ut.ee/). The amplified products were sequenced and analyzed with CRISPResso2 software(https://github.com/pinellolab/CRISPResso2).

Cas-OFFinder (http://www.rgenome.net/cas-offinder/) was used to predict off-targets of PCSK9-sg1, allowing for up to 3 mismatch bases and up to 1 DNA or RNA bulge. Following analysis and consolidation of the 54 search information, a total of 28 sites were identified, comprising 1 on-target site, 3 off-target sites with 3 base mismatches, 22 off-target sites with 2 base mismatches and 1 RNA bulge, and 2 off-target sites with 2 base mismatches and 1 DNA bulge.

### R-loop Assay

To assess guide RNA-independent DNA off-target effects of the base editor, HEK293T cells were seeded at a density of 70,000 cells per well in a 48-well plate 24 hours before transfection. Cells were co-transfected using PEI with 250 ng of base editor plasmid, 250 ng of sgRNA plasmid, 250 ng of nSaCas9 plasmid, and 250 ng of nSaCas9 sgRNA plasmid. 72 hours after transfection, the edited genome was extracted, the target sequence was amplified and sequenced, and the editing results were quantified using the methods described above.

### mRNA Preparation

All the ABE mRNA variants were synthesized by in vitro transcription and purified using the RNasy Mini Kit (Qiagen, 74143) in accordance with the manufacturer’s instructions. For the in vitro editing experiments in cell lines and primary cells, linearized plasmids containing the full-length ABE coding sequences were utilized as templates, These were further transcribed by the HiScribe® T7 High Yield RNA Synthesis Kit (NEB, E2040S) and followed by polyadenylation with the Poly(A) Tailing Kit (Invitrogen, AM1350). For the in vivo editing experiments in monkeys, the construct was modified by inserting a 120nt-polyA tail following the 3’ untranslated region sequence (3’UTR). The modified mRNA was generated by substituting the UTP with N1-methylpseudouridine, which was further quantified using a NanoDrop spectrometer and analyzed via a Bioanalyzer (Agilent). Chemically modified guide RNAs, featuring 2’-O-methyl and 3’ phosphorothioate modifications at the first and last three nucleotides, were ordered from GenScript.

### LNP formulation

For cynomolgus monkey and cellular studies, LNPs were formulated as previously described29,30, with the lipid components (ionizable lipid, 1,2-distearoyl-sn-glycero-3-phosphocholine, cholesterol and a PEG-lipid) being rapidly mixed with an aqueous buffer solution containing hpABE5.20 mRNA and PCSK9 sgRNA in a 1:1 ratio by weight. The LNPs had a particle size of 70-90 nm, with a polydispersity index of <0.2 as determined by dynamic light scattering (Malvern NanoZS Zetasizer) and >90% total RNA encapsulation as measured by the Quant-iT Ribogreen Assay (Thermo Fisher). The resulting LNP formulations were subsequently dialysed against 1× PBS and filtered using a 0.2-μm sterile filter. The LNPs had an average diameter of 70-90 nm, with a polydispersity index of <0.2 as determined by dynamic light scattering and >90% total RNA encapsulation as measured by the Quant-iT Ribogreen Assay.

### LNP treatment of cynomolgus monkeys

The cynomolgus monkey studies were approved by the ECNU Animal Care and Use Committee and in direct accordance with the Ministry of Science and Technology of the People’s Republic of China on Animal Care guidelines. The cynomolgus monkey study was performed at Wincon TheraCells Biotechnologies Co.,Ltd using male cynomolgus monkeys. The monkeys were 3-4 years of age and 3-5 kg in weight at the time of study initiation. All monkeys were genotyped at the PCSK9 editing site to ensure that any monkeys that received the ABE8.8 and PCSK9-1 LNPs were homozygous for the protospacer DNA sequence perfectly matching the gRNA sequence; otherwise, monkeys were randomly assigned to various experimental groups, with collection and analysis of data performed in a blinded fashion. The sample sizes for the experimental groups were chosen based on ethical principles (that is, the minimum necessary number of monkeys). The monkeys were premedicated with 1 mg kg−1 dexamethasone, 0.5 mg kg−1 famotidine and 5 mg kg−1 diphenhydramine on the day before LNP administration and then 30–60 min before LNP administration. The LNPs were administered via intravenous infusion into a peripheral vein and the infusion speed is 0.5mL/min. Control monkeys that received PBS instead of LNPs experienced the same infusion conditions. For blood chemistry samples, monkeys were fasted overnight before collection via peripheral venipuncture. In all cynomolgus monkey studies, samples were typically collected on the following schedule: day –10, day –7, day –5, day 1 (6 h after LNP infusion), day 2, day 3, day 8, day 15, day 22 and day 29. In the long-term study, samples were also collected at day 21 and day 28 and have generally been collected every month thereafter. Blood samples were analysed by the study site for LDL cholesterol, HDL cholesterol, total cholesterol, triglycerides, AST, ALT, alkaline phosphatase, γ-glutamyltransferase, total bilirubin and albumin. For each analyte, the baseline value was calculated as the mean of the values at day –10, day –7 and day –5. A portion of each blood sample was sent to the investigators for PCSK9 measurement using the LEGEND MAX Human PCSK9 ELISA Kit (BioLegend), with recombinant cynomolgus monkey PCSK9 (PC9-C5223, Acro) for standardization. In the long-term cynomolgus monkey study, each monkey underwent an ultrasonography-guided percutaneous liver biopsy using a 16-gauge biopsy needle, performed under general anaesthesia, on day 15 and day 29. The samples were processed with the Tiangen DP304 kit to isolate genomic DNA.

Data availability

All RNA sequencing data generated in this article can be found at the National Center for Biotechnology Information’s Sequence Read Archive with accession code PRJNAxxxxxx. All data supporting the findings of this study are available within the article and Supplementary Information files and also are available from the corresponding author upon request. Source data are provided with this paper.

## Acknowledgements

This work was supported by the National Key R&D Program of China 2023YFC3403401 & 2024YFA1803301 (Y.W.), the Shanghai Agricultural Science and Technology Innovation Program K2023001 (Y.L.), the National Natural Science Foundation of China 82270125 (Y.W.), 32300667 (J.L.), 32371535 (S.C.), 82450107(D.W.), the project of Shanghai Municipal Science and Technology Commission 23HC1400400 (Y.W.), the National Program for Support of Top-Notch Young Professionals (Y.W.). The flow cytometry, qPCR, RP-HPLC were performed at the Instruments Sharing Platform of School of Life Sciences, East China Normal University. The authors thank Ying Zhang for help with flow cytometry.

## Author information

These authors contributed equally: Jiaoyang Liao, Hongling Zhang.

These authors jointly supervised this work: Zijun Wang, Yuxuan Wu, Yuming Lu.

## Authors and Affiliations

Shanghai Frontiers Science Center of Genome Editing and Cell Therapy, Shanghai Key Laboratory of Regulatory Biology, Institute of Biomedical Sciences and School of Life Sciences, East China Normal University, Shanghai, China

Jiaoyang Liao, Shuanghong Chen, Shenlin Hsiao, Yanhong Jiang & Yuxuan Wu

YolTech Therapeutics, Shanghai, China.

Hongling Zhang, Zijun Wang, Wendan Ren, Changrui Feng, Da Xie, Chongping Lai, Wenlong Wang, Yanqiu Zheng, Wenhui Cai, Yuming Lu & Yuxuan Wu

School of Agriculture and Biology, Shanghai Jiao Tong University, Shanghai, China. Yuming Lu

Shenzhen Branch, Guangdong Laboratory for Lingnan Modern Agriculture, Key Laboratory of Synthetic Biology, Ministry of Agriculture and Rural Affairs, Agricultural Genomics Institute at Shenzhen Chinese Academy of Agricultural Sciences, Shenzhen, China

Hu Feng & Erwei Zuo

Shanghai Institute of Hematology, State Key Laboratory of Medical Genomics, National Research Center for Translational Medicine at Shanghai, Ruijin Hospital Affiliated to Shanghai Jiao Tong University School of Medicine, Shanghai, China

Dawei Wang

## Contributions

J.L., H.Z., Z.W., Y.L. and Y.W. completed the conceptualization of the study. J.L., H.Z., S.C. and S.H. designed and performed experiments. J.L., H.F. and C.F. performed computational analyses, oversaw data analysis/interpretation, under the supervisition of E.Z. and Y.W.. Z.W. developed a novel LNP-mRNA delivery system and, with the assistance of W.R., D.X. and Y.J., carried out the production of LNP-mRNA and its delivery to both cells and animas. C.L., Y.Z., W.C. D.W. and W.W. helped with the molecular and cellular experiments. J.L. wrote the manuscript with input from all authors. E.Z., Z.W., Y.L. and Y.W. contributed to the final version of manuscript.

## Competing interests

H.Z., C.L., and Z.W. have filed and been granted patent applications related to this work. Z.W., Y.L. and Y.W. are co-founders of YolTech Therapeutics, Shanghai, China. The other authors declare no competing interests.

