## Supplemental information for "Sequential Amino Acid Mutagenesis-Driven De Novo Evolution of Adenine Deaminases Enables Efficient In Vivo Base Editing in Primate"

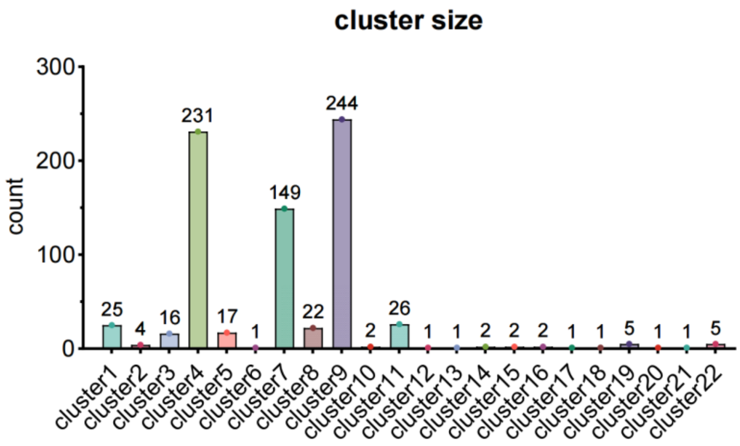


**Figure S1 |** The number of members in each cluster is plotted in a bar chart after clustering the 759 candidate adenine deaminase protein structures. The corresponding number is marked at the top of each bar; for example, cluster 1 contains a total of 25 members, including ec_tadA.

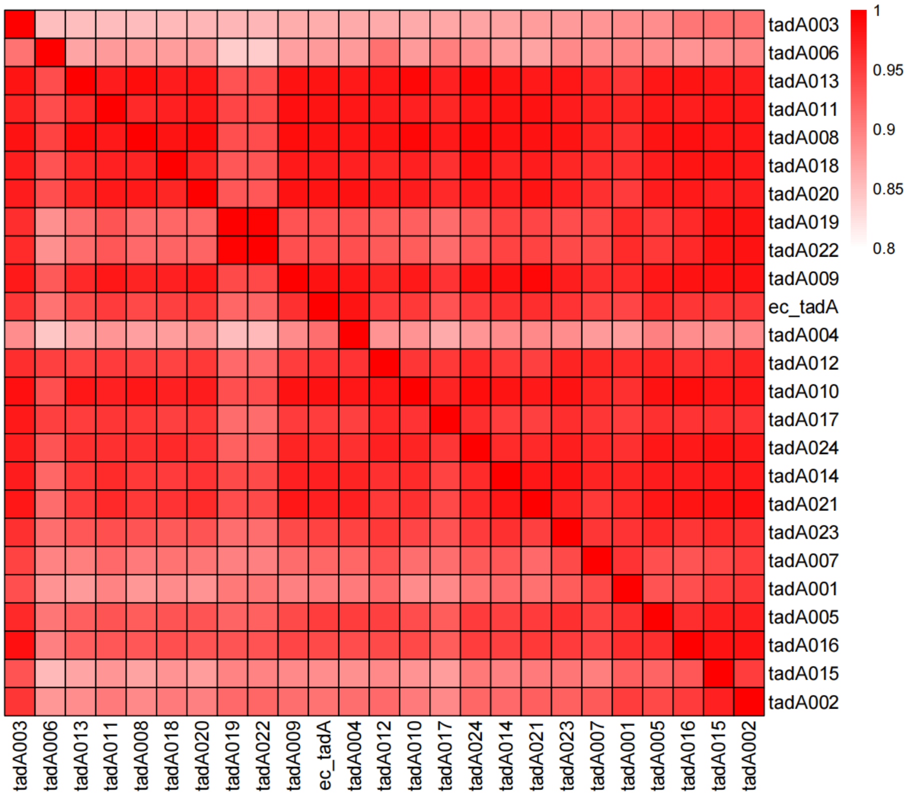


**Figure S2 |** Heat map of structural similarity scores between members of cluster 1. The results indicate that the structural similarity among the components in cluster 1 is very high, with scores exceeding 0.8.

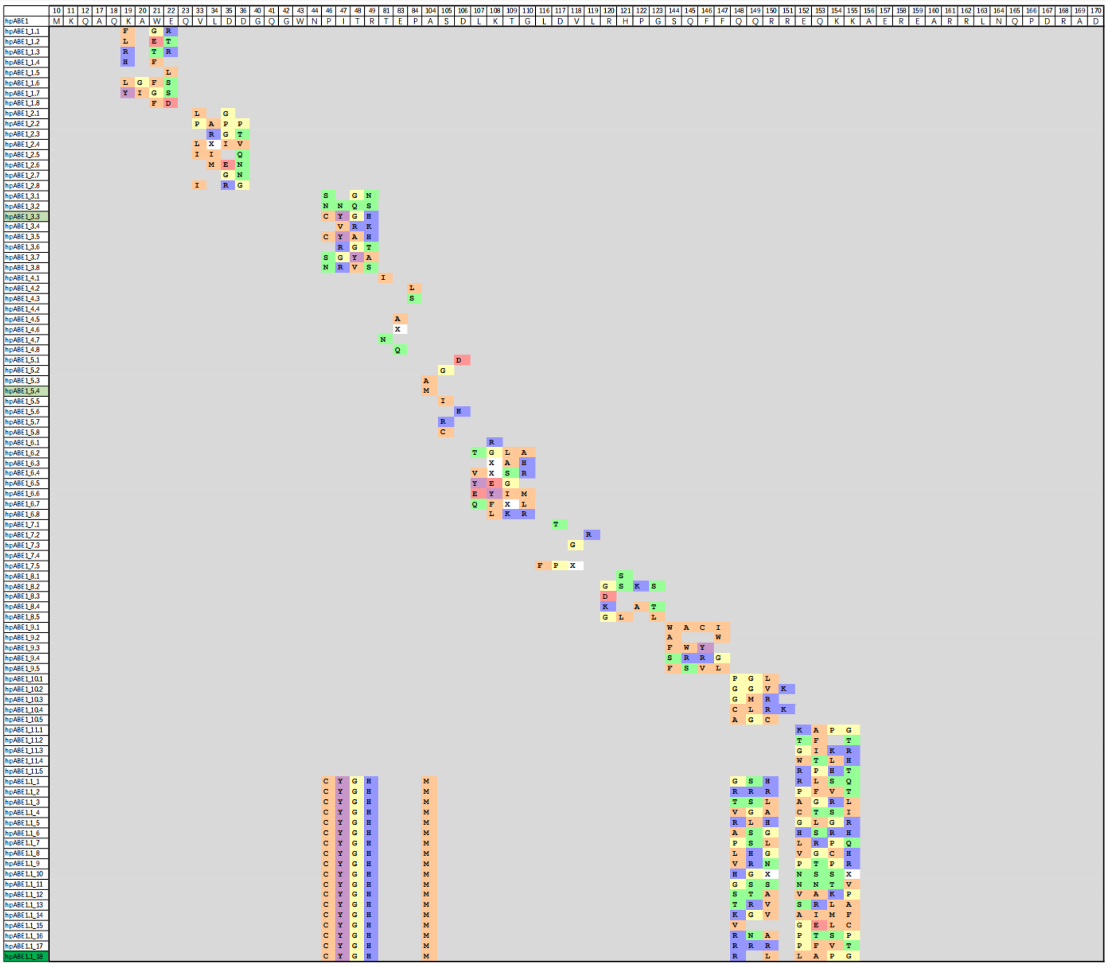

**Figure S3 |** Amino acid profiles of hpTadA variants identified during round1 directed evolution. These profiles are based on the hpTadA1 variant, which was derived from hpTadA0 with 17 point mutations (corresponding to the 17 mutations in tadA8e relative to wild-type ecTadA). The variant names are highlighted in light green for the regional dominant variants hpABE1_3.3 and hpABE1_5.4, and in dark green for the optimal variant from this round of evolution, hpABE1.1_18. The individual letters represent the various amino acids, colored according to the Zappo color scheme to distinguish differences in hydrophobicity, polarity, and charge.

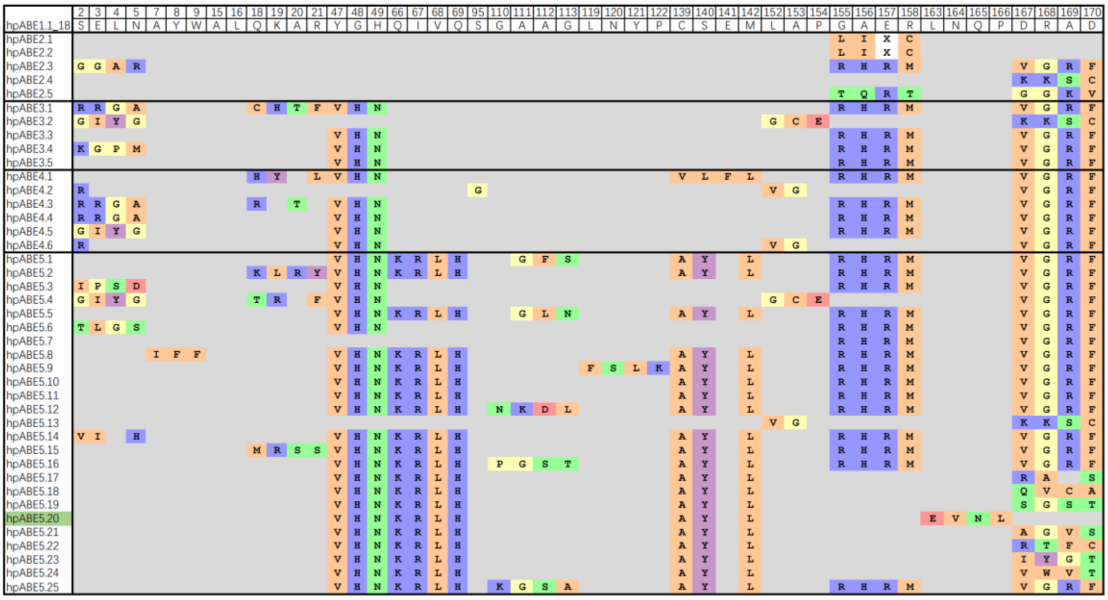

**Figure S4 |** Amino acid changes in hpTadA variants identified during rounds 2-5 of directed evolution. The amino acid changes in these variants are compared to the hpTadA1.1_18 variant. The best variant from round 5, hpABE5.20, is highlighted in light green. The individual letters represent various amino acids, colored according to the Zappo color scheme to distinguish differences in hydrophobicity, polarity, and charge.
